# Discovery of a novel MarR-type transcriptional regulator that controls cell death in *Bacillus subtilis* biofilms

**DOI:** 10.1101/2023.11.22.568295

**Authors:** Gillian McClennen, Leticia Lima Angelini, Gabriel Fox, Veronica Godoy-Carter, Yunrong Chai

**Author notes:** Both authors contributed equally to this work (Gillian McClennen, Leticia Lima Angelini).

## Abstract

Pulcherriminic acid (PA) is a cyclic-_L_-leu-_L_-leu di-peptide produced by *Bacillus subtilis* during biofilm formation. When secreted, PA strongly chelates extracellular iron and forms a reddish-brown pigment, pulcherrimin. Production of pulcherriminic acid and formation of pulcherrimin modulate iron homeostasis in *B. subtilis*. Pulcherriminic acid also functions as an antioxidant to protect cells from increasing oxidative stress during biofilm formation. We previously showed that PA is involved in gene regulation, differentially regulating hundreds of genes in *B. subtilis*. One of the strongly upregulated genes by PA is *yhjH*, encoding a putative MarR-type transcription repressor. In this study, we characterized the regulation of the *yhjH* gene by PA, by PchR, a known transcription repressor for PA biosynthesis, and by YhjH itself. We also found that high expression of *yhjH* triggers rapid cell lysis in *B. subtilis*. Results from RNA-seq suggest that YhjH differentially regulates about 180 genes, among which there is a significant number of prophage genes. Lastly, we propose that YhjH be re-named as PcdR, for “<u>p</u>ulcherriminic acid <u>c</u>ell <u>d</u>eath <u>r</u>egulator”.

## Introduction

Biofilms are intricate macroscopic structures formed by communities of bacteria. These structures are composed of a complex matrix of polysaccharides, proteins, and nucleic acids (3). The biofilm matrix provides protection for bacteria within biofilms and facilitates an environment where bacterial cells adopt specialized roles and share information. Protection provided by biofilms protects pathogenic bacteria from antimicrobial agents, which poses concerns for clinical settings and the food industry (4). Understanding biofilm development and maintenance presents an opportunity to find new solutions to prevent or treat biofilm-associated infections.

Bacteria communicate through intercellular communication pathways (e.g. quorum-sensing) to express virulence factors, spread antibiotic resistance, and signal shifts from planktonic to biofilm lifestyles (5). The formation, maintenance, and disassembly of biofilms relies on this cell-cell communication. As epicenters of bacterial activity, with a rich array of molecules that direct microbial life, biofilms provide an interesting system to study cell-cell communication.

Pulcherrimin is a reddish-brown, iron-chelating pigment that is synthesized during biofilm formation by *Bacillus subtilis* (4). The pigment is a ferric salt of pulcherriminic acid (PA), which is a cyclic di-leucine peptide product that is synthesized from two charged leucyl-tRNA molecules (6, 7). PA is synthesized inside the cell and secreted to bind extracellular ferric iron (Fe^3+^) and form the pigmented pulcherrimin (Figure 1) (8). In some yeast species, pulcherrimin has been found to exhibit antimicrobial properties against plant pathogens, including microscopic fungi and other bacteria (9). However, the biological role of the pigment in *B. subtilis* is less understood. One study showed that the strong chelation of iron by PA causes growth arrest during *B. subtilis* biofilm formation (8, 10). Charron-Lamoureux, as well as our group, also showed that pulcherrimin functions as an antioxidant to protect cells from increasing oxidative stress they experience under biofilm conditions (11, 12).

**Figure 1:**
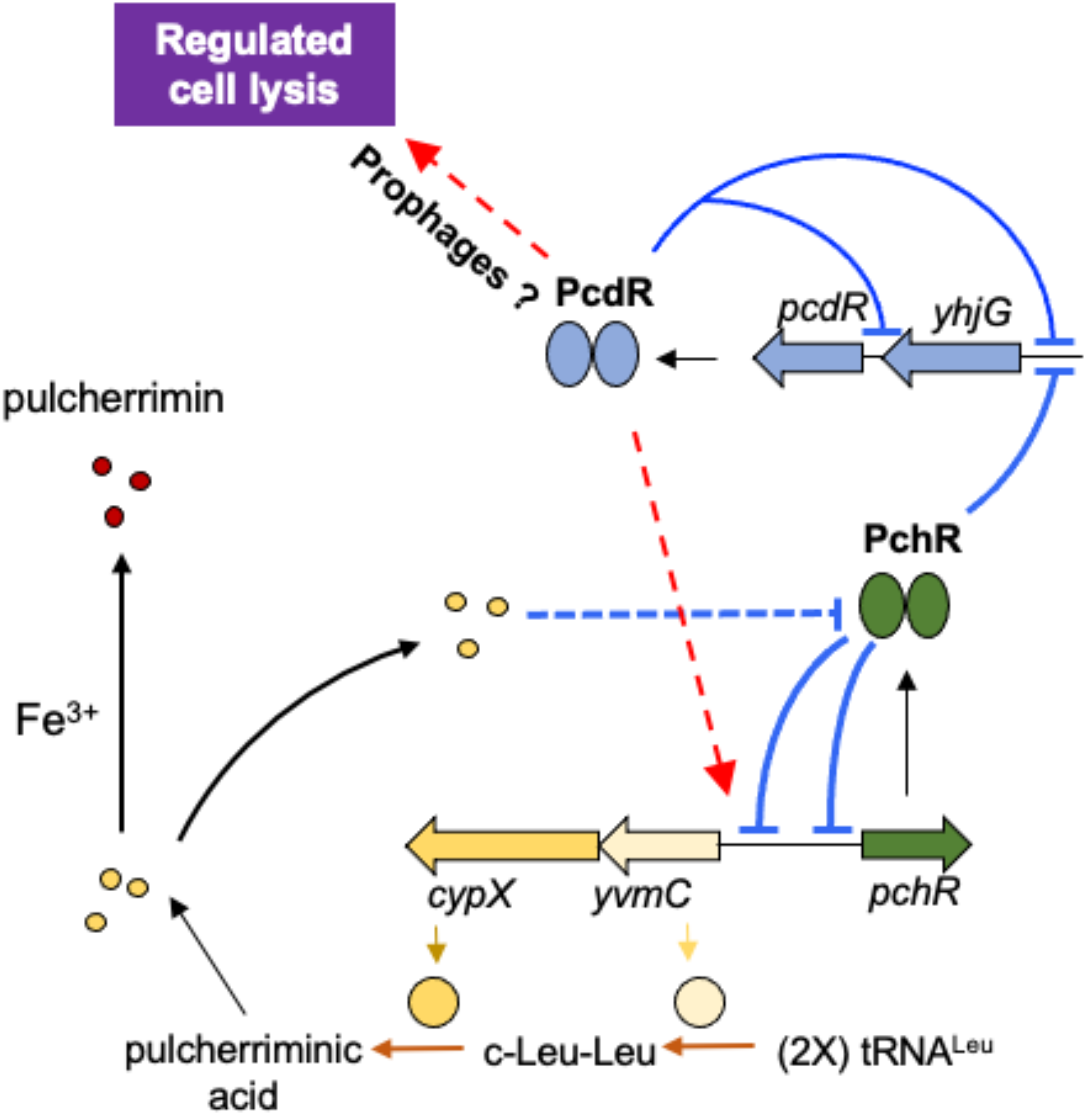
A working model of pulcherriminic acid (PA) production and *yhjH* regulation. PA is synthesized by the cyclodipeptide synthase YvmC, which cyclizes two leucine residues into cyclo-di-leucine (c-Leu-Leu), and the cytochrome oxidase CypX, which oxidizes c-Leu-Leu to form PA. After secretion, PA can either bind ferric iron to form pulcherrimin, or mediate a putative signal transduction pathway to antagonize the action of PchR, the repressor for PA biosynthetic genes. PchR also represses the operon of *yhjG-pcdR*. PcdR represses both the *yhjG-pcdR* operon and a promoter located in the intergenic region between *pcdR* and *yhjG*. In addition, PcdR positively regulates PA production and its overexpression triggers a cell lysis phenotype.

Cyclic dipeptides can act as modulators of microbial behavior in response to the environment. The study of the role of dipeptides as signaling molecules has implications for gut microbiome modulation of the immune system, gut-brain interactions, control of pathogen virulence, biofilms, and quorum sensing (13, 14). One recent example was the characterization of cyclo- Val-Leu as a quorum-sensing molecule with the ability to stimulate growth and promote biofilm formation of the gut commensal *Bifidobacteria* (15, 16). Recent interest in this field led us to take another look at the conventional view of pulcherriminic acid (PA) as simply a pulcherrimin precursor. Our previous study has suggested that PA is also involved in gene regulation in *B. subtilis*; its presence differentially regulates hundreds of genes involved in biofilm development, sporulation, stress response, surface polysaccharide biosynthesis, and iron homeostasis (12).

Given that PA is a cyclic-di-peptide derivative produced under biofilm conditions, we hypothesized that this secondary metabolite functions as a signaling molecule that directly modulates biofilm development by a yet-to-be characterized signal transduction mechanism.

Programmed cell death is the process of genetically controlled suicide behavior in bacterial populations (17). Although bacteria are individual cells, they work together in a multicellular community while in biofilms. This process, therefore, requires cells to behave altruistically. Programmed cell death is used by bacteria to contribute extracellular DNA (eDNA) and proteins to the biofilm matrix and create wrinkled and fruiting body phenotypes on a macroscopic scale (18–20). There are a few different mechanisms by which bacteria achieve regulated cell death. In *Pseudomonas aeruginosa*, cell death phenotypes were caused by the activation of dormant prophages in the genome (21). Bacteria have also co-opted phage-mediated cell lysis with murein hydrolases, like the *cid* and *lrg* operons in *Staphylococcus aureus*, which are analogous to phage holin/antiholin systems (17). Other mechanisms for programmed cell death involve autolysins, which hydrolyze the peptidoglycan cell wall, causing the bacterial cell to burst due to osmotic pressure (19). In *B. subtilis*, localized cell death contributes to wrinkle formation during biofilm formation, yet the molecular mechanism for the observed cell death is not well understood (18).

The aim of this study was to characterize YhjH: a novel MarR-type transcriptional regulator whose gene expression is activated by pulcherriminic acid (PA) during *B. subtilis* biofilm formation (12, 22). We found that high expression of *yhjH* causes an extreme cell lysis phenotype that likely plays a role in programmed cell death in *B. subtilis* biofilm development. We identified that YhjH represses a binding motif present in the gene promoter for the operon of *yhjG* and *yhjH* and in a short intergenic promoter between these two genes. Furthermore, based on global transcriptome analysis and binding motif search, we propose that this cell lysis phenotype observed in *B. subtilis* can be genetically programmed through the involvement of upregulation of prophage genes induced under the control of YhjH. YhjH is currently annotated as a MarR-type transcriptional regulator, but its function was unknown. From this study, we propose that YhjH be renamed PcdR (<u>P</u>ulcherriminic acid <u>c</u>ell <u>d</u>eath <u>R</u>egulator), given its clear induction of cell death after regulation by pulcherriminic acid.

## Results

### The *pcdR* gene is activated by pulcherriminic acid

We performed an RNA-seq experiment between the wild-type strain 3610 and a strain lacking the pulcherriminic acid (PA) biosynthetic genes (LA10, Δ*yvmC-cypX*) to compare the global transcriptomes and to study the effects of PA biosynthetic gene deletion on *B. subtilis* biofilm development (12). 692 genes are differentially regulated when using log2FC of ±1 as the cutoff. Among the significantly downregulated genes in the PA deficient strain (Δ*yvmC-cypX*) are *pchR* and *pcdR* (log2FC of -4.8 and -2.0 for *pchR* and *pcdR*, respectively), encoding two MarR-type transcription repressors (12). PchR is a known transcription repressor for the Δ*yvmC-cypX* operon, while PcdR (formerly known as YhjH) is a putative MarR-type transcription regulator of unknown function (23, 24). Interestingly, PchR and PcdR share ∼42% identity in their amino acid sequences (Supplementary Figure 1). In addition, *yhjG*, the gene next to *pcdR* in the apparent *yhjG-pcdR* operon, is also downregulated in the Δ*yvmC-cypX* strain (log2FC of -2.29) (12, 24).

To confirm that the *yhjG-pcdR* genes are activated by PA, the promoter of the *yhjG-pcdR* operon was fused to the gene for green fluorescent protein (GFP) and the reporter fusion (P*_yhjG_*-*gfp*) was introduced into the wild-type strain, the Δ*yvmC-cypX* strain, and the complementation strain (LA33, Δ*yvmC*-*cypX*, *amyE*::P*_hyspank_*-*yvmC-cypX*). Activities of the promoter fusion were determined by assessing the fluorescence intensity of the cells expressing P*_yhjG_-gfp* using the ImageJ script from Brychcy, et al (2). It was found that the activity of P*_yhjG_-gfp* was significantly decreased in the Δ*yvmC-cypX* strain compared to the wild-type, while complementation of Δ*yvmC-cypX* increased the activity of the promoter fusion to above wild-type levels (Figure 2A- B). Similar results were obtained when the *yhjG-pcdR* promoter was fused to *lacZ*; expression of P*_yhjG_*-*lacZ* was significantly higher in the wild-type than in the Δ*yvmC*-*cypX* strain (Figure 2C). Based on these results, we conclude that PA production during biofilm formation activates the *yhjG-pcdR* genes.

**Figure 2.**
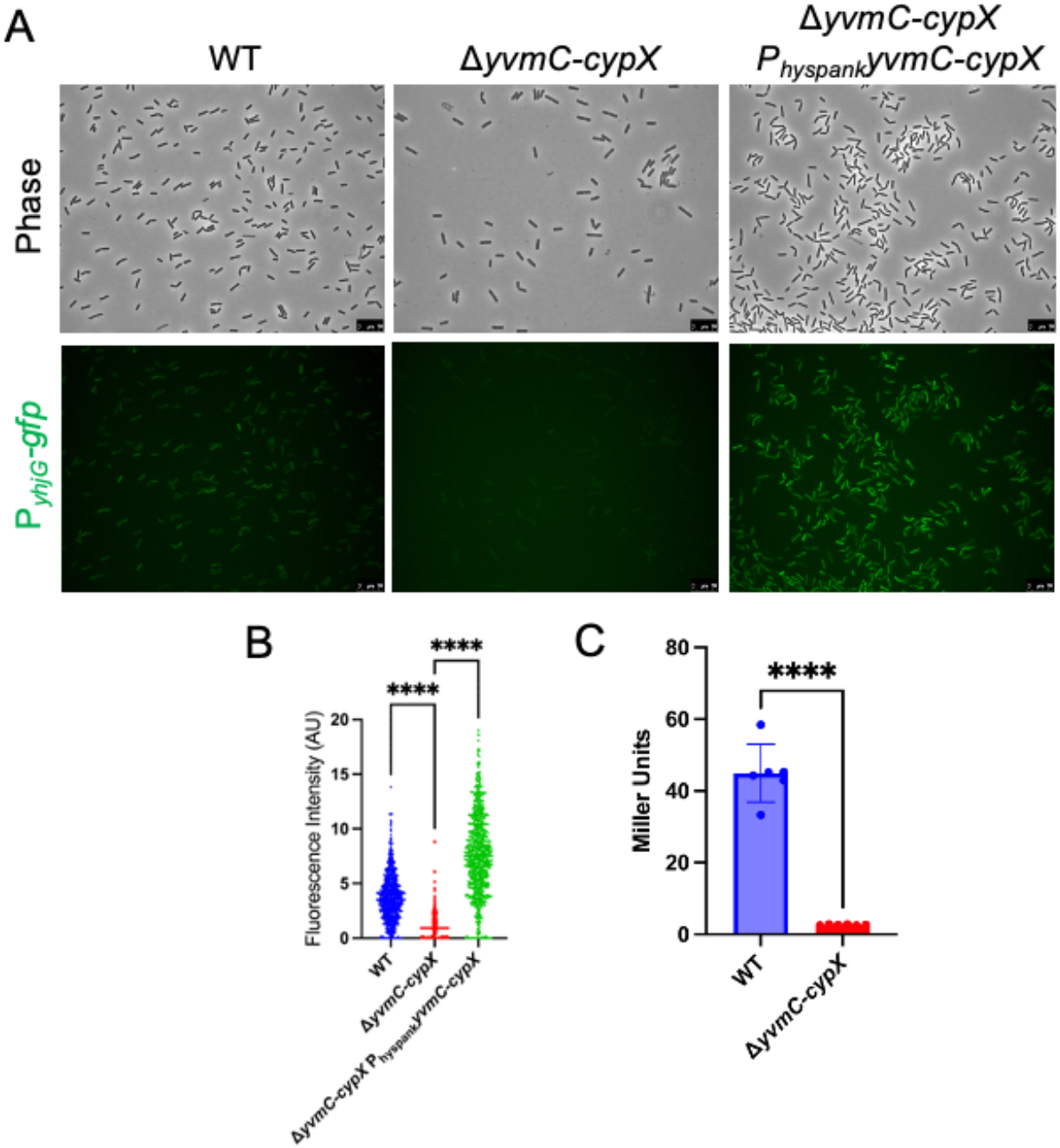
The promoter of *yhjG-pcdR* is activated in the presence of pulcherriminic acid. (A) The promoter of the *yhjG-pcdR* operon was fused to *gfp* and introduced into the wild-type (3610), the *yvmCcypX* deletion mutant, and the complementation strain. Pellicle biofilms by the above strains were developed for 3 days before cells were harvested and imaged by fluorescence microscopy. Scale bar, 10 μm (applies to all panels). **(B)** Fluorescence density of cells was quantified by ImageJ using a modified script outlined in Brychcy, et al. (2).Significance was calculated with Wilcoxon’s T-test using Graphpad Prism. **(C)** The promoter of *yhjG-pcdR* was also fused to *lacZ* and the fusion was introduced into the wild type and the *yvmC-cypX* deletion mutant. Assays of β-galactosidase activities were then conducted for the above engineered reporter strains and were repeated in biological duplicate. **** represents p <0.0001.

### The *pcdR* gene is both repressed by PchR and autorepressed by PcdR

PchR is a known transcriptional repressor for the PA biosynthetic genes as well as itself (23). Interestingly, when activities of P*_yhjG_-gfp* were assayed in the wild type and the Δ*pchR* deletion strain, we found that the *pchR* gene deletion greatly increases the expression of *yhjG-pcdR* (Figure 3), suggesting that in addition to the PA biosynthetic genes, PchR also represses *yhjG- pcdR*.

**Figure 3.**
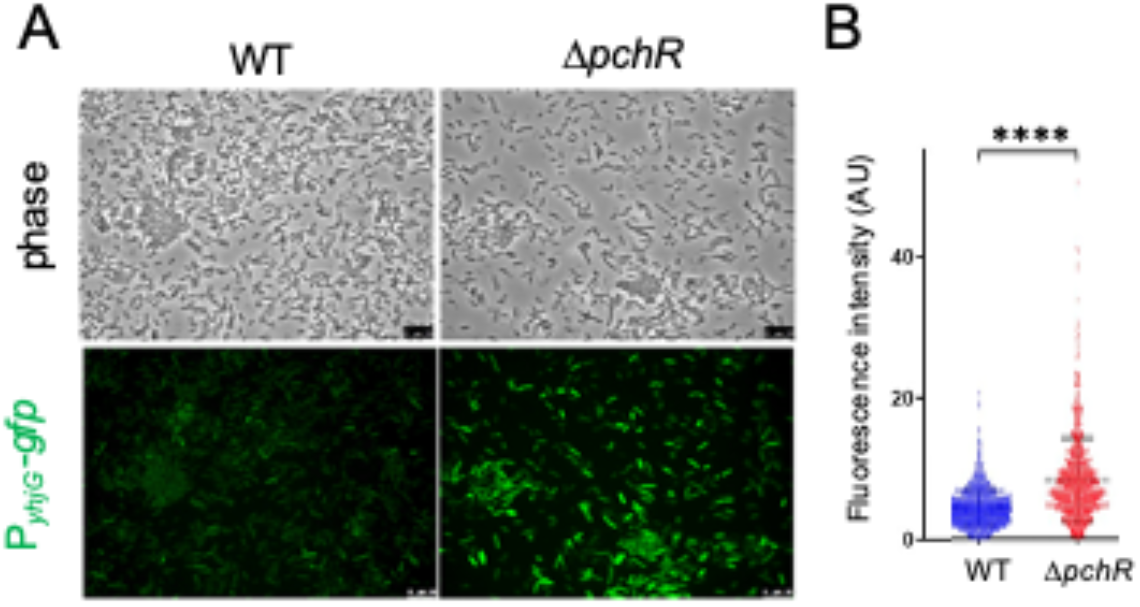
PchR represses the *yhjG-pcdR* operon. (A) The P*yhjG*-*gfp* promoter fusion was introduced into the wild-type and the Δ*pchR* mutant background. Cultures were grown for 3 days in pellicle biofilm conditions before cells were harvested and imaged by fluorescence microscopy. Scale bar, 10 μm (applies to all panels). (**B**) Fluorescence was calculated using ImageJ and plotted with Prism (1). **** indicates p value <0.0001.

Like PchR, PcdR is a MarR-type transcriptional regulator (24, 25). We then tested whether PcdR autoregulates itself, similar to what was seen in PchR (23). To do so, the P*_yhjG_-gfp* promoter fusion was introduced into the Δ*pcdR* deletion strain, and its activity was compared between the wild-type and the Δ*pcdR* strain. It was found that the activity of P*_yhjG_-gfp* was significantly higher in the Δ*pcdR* strain (Figure 4A-B). Interestingly, not only is the promoter of the *yhjG- pcdR* operon repressed by PcdR, PcdR also represses an internal promoter located in the short intergenic region (76-bp) between *yhjG* and *pcdR* (Figure 1). When we fused the intergenic sequence between *yhjG* and *pcdR* with the *lacZ* gene and introduced the resulting reporter (P*_pcdR_*- *lacZ*) into both the wild-type and the Δ*pcdR* strain, the reporter showed a significantly higher activity in the Δ*pcdR* strain than in the wild-type strain (Figure 4C). Our results thus suggest that PcdR represses its own gene from two independent promoters, and that the *yhjG-pcdR* genes are also repressed by PchR. How PA activates *yhjG-pcdR* is not known. We suspect that PA antagonizes PchR-mediated repression on *yhjG-pcdR* through a yet-to-be characterized mechanism (Figure 1; LA, unpublished data).

**Figure 4.**
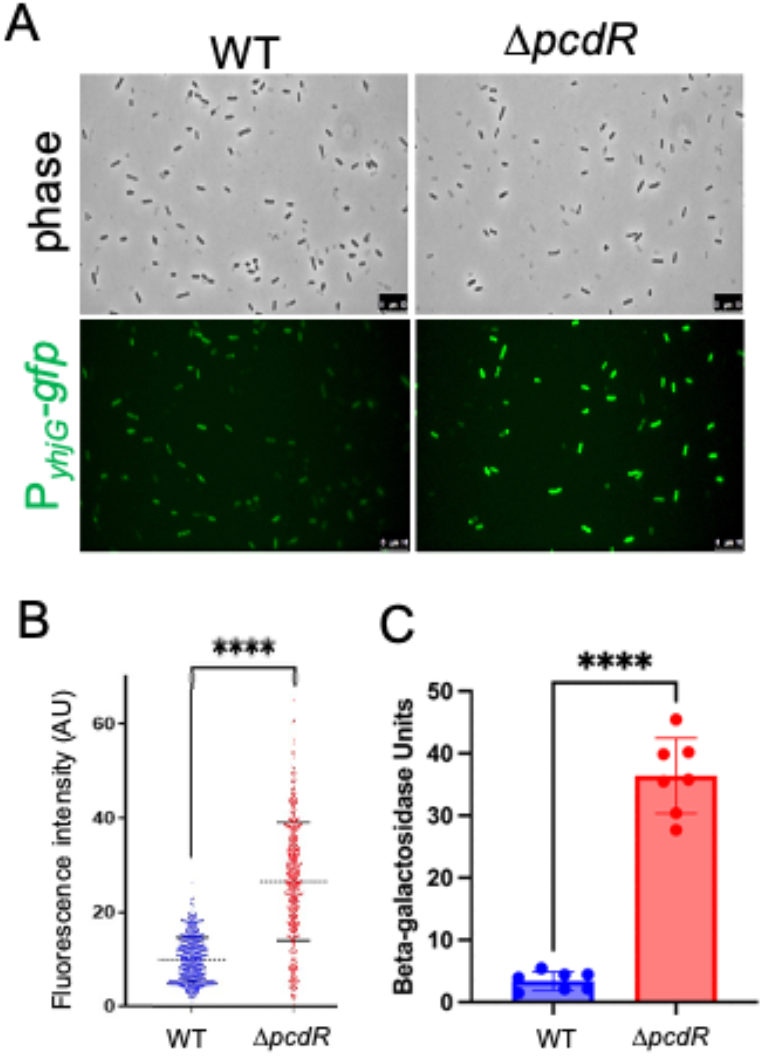
PcdR autorepresses its own gene. (A) Samples were harvested after 3 days of growth in pellicle biofilms. P*yhjG-gfp* expressing cells were lightly sonicated to break up bundled chains and associated biofilm matrix and imaged by fluorescence microscopy. Scale bar, 10 μm (applies to all panels). **(B)** Fluorescence intensity was calculated using MicrobeJ (1). **(C)** Similarly, P*pcdR*-*lacZ*expressing cells were mildly sonicated and β-galactosidase assays were performed. **** indicates p value < 0.0001.

### Loss of *pcdR* attenuates pulcherrimin production while its overexpression leads to rapid cell lysis

To investigate the function of PcdR, we constructed a /-*pcdR* deletion mutant. Interestingly, the /-*pcdR* mutant exhibits impaired pigment production as shown in colony biofilms (Figure 5A). Our preliminary result thus suggests that PcdR is a novel regulator for PA biosynthesis, but opposite to PchR. This regulation is likely indirect since PcdR is predicted to be a transcription repressor. Our results also imply that PA biosynthesis could be regulated by a feedback mechanism; PA production leads to *pcdR* expression and *pcdR* expression results in even higher PA production. In addition, we found that pigment production in the double mutant of /-*pcdR*/-*pchR* moderated when compared to that in the /-*pchR* single mutant, but higher than that in the *pcdR* single mutant (Figure 5B). This result indicates that PchR and PcdR could regulate PA biosynthesis independently.

**Figure 5:**
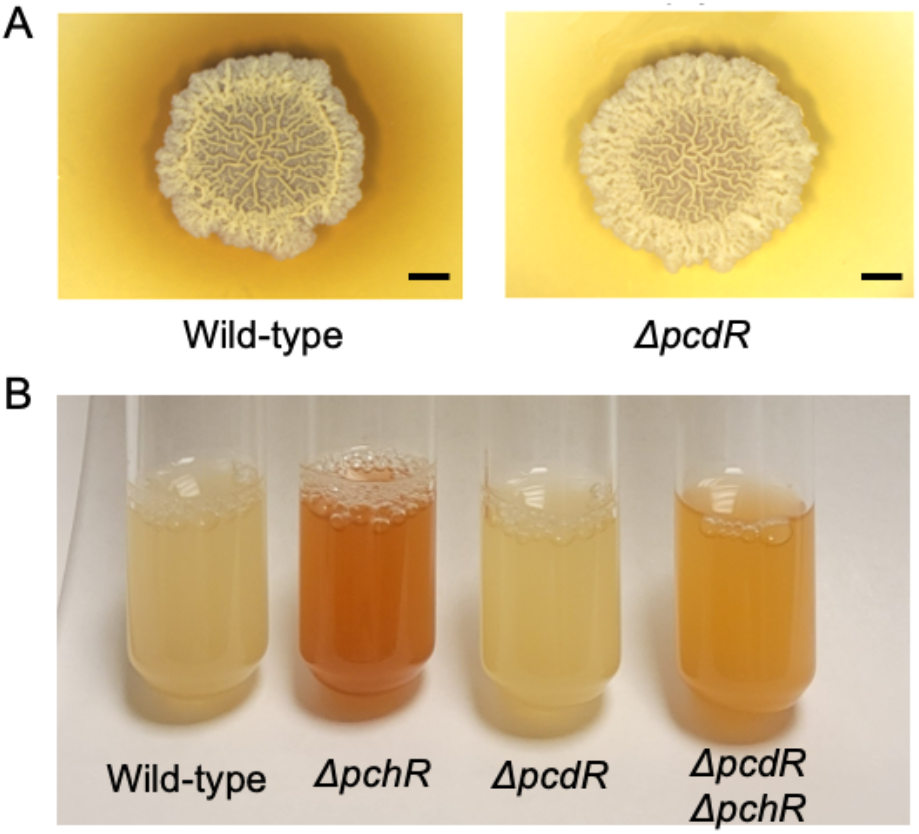
The *pcdR* gene deletion decreases pulcherrimin production. (A) Colony biofilms formed by the wild-type strain and the Δ*pcdR* deletion strain were plated and visualized after 3 days of incubation. The Δ*pcdR* strain exhibited less pulcherrimin production than the wild type. Scale bar, 250 mm. **(B)** Cultures were grown overnight in LBGM supplemented with 100 μM of ferric iron and color of the cultures was visualized the next morning. Pulcherrimin overproduction in a pulcherrimin repressor knockout strain (Δ*pchR*) was mitigated in a *pcdR* knockout strain (Δ*pchR* Δ*pcdR*).

To further our investigation on PcdR, a strain was engineered to have a second copy of *pcdR* fused to an IPTG-inducible promoter (P*_hyspank_*-*pcdR*) and the construct was integrated at the *amyE* locus on the chromosome. This engineered strain overexpresses *pcdR* upon IPTG induction.

Initially, we wanted to test if overexpression of *pcdR* leads to hyperproduction of pulcherrimin. However, it was quickly found that overexpression of *pcdR* induced rapid cell lysis. The growth of the engineered strain was tested at varying concentrations of IPTG, and it was found that cells exhibited rapid lysis at an IPTG concentrations higher than 25 µM, , whereas no death phenotype was observed at a final concentration of 5 µM or lower (Figure 6A).

**Figure 6.**
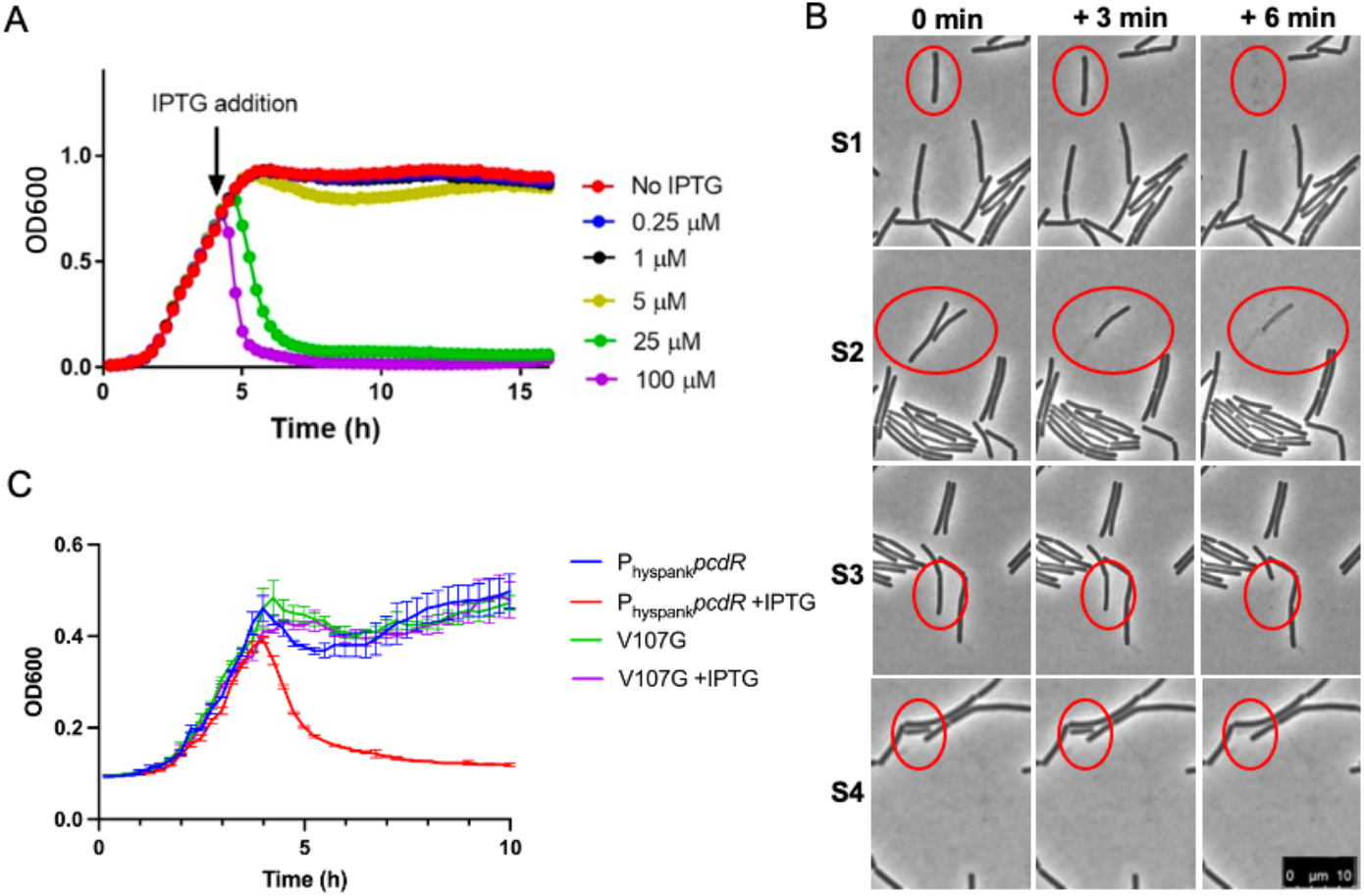
Overexpression of *pcdR* causes a rapid cell lysis phenotype. (**A)** An IPTG-inducible *pcdR* was constructed in the wild-type *B. subtilis* background (*Physpank-pcdR*) and grown under shaking conditions at 37°C with varying concentrations of IPTG. (**B)** Cells bearing P*hyspank-pcdR* were grown to exponential phase and inoculated onto an agar pad with 25 μM IPTG. Confocal microscopy images were taken every 3 minutes. Shown are frames of time lapse video taken 3 minutes apart. Visible cells slightly shrink (second panel) then completely disappear within a 3 min time span (third panel). Cells of interest are circled in red. Scale bar, 10 μm (applies to all panels) **(C)** Overexpression of the *pcdR* construct in Suppressor 8, a strain carrying the V107G PcdR suppressor mutation in the overexpression construct, does not exhibit the rapid death phenotype after IPTG induction.

A time-lapse microscopic imaging was performed to observe cell death in real-time and to characterize the dynamics of cell death. Cells were inoculated onto an agar pad supplemented with 25 µM IPTG. Snapshot images were taken every 3 minutes. We observed a rapid cell “pop” phenotype: individual cells disappeared within minutes (highlighted by red circles, Figure 6B). No clear morphological deformation was observed for cells undergoing lysis. We thus conclude that high expression of *pcdR* triggers rapid cell lysis in *B. subtilis*.

### Suppressor mutants emerge after prolonged incubation post cell lysis

Upon continued incubation of the cultures after complete cell lysis, we observed turbidity in selected test tubes the following morning, suggesting emergence of suppressor mutants. We purified selected putative suppressor mutants to single colonies and verified that the suppressor mutants are no longer sensitive to PcdR-mediated cell lysis when inoculated into fresh media supplemented with IPTG. To characterize genes involved in this novel cell death phenotype, we performed whole genome sequencing for 8 selected suppressor mutants. Not surprisingly, most of these suppressor mutants developed mutations in the *pcdR* overexpression construct itself, suggesting that it is indeed the overexpression of PcdR that causes the phenotype. One of the suppressors was different; it has a missense mutation in the *pcdR* coding sequence that results in a V107G change in amino acid sequence (Supplementary Figure 1). This mutation occurs in the DNA-binding wing of the PcdR protein, as predicted by Alphafold and annotated by Deochand and Grove (26). This mutation likely abolishes PcdR DNA binding ability. Remarkably, the mutation does eliminate the death phenotype (Figure 6C). This result reinforces the idea that the DNA-binding activity of PcdR is responsible for the death phenotype.

### Identification of YhjH regulon by RNA-seq

To explore the mechanism of cell lysis mediated by PcdR, we performed an RNA-seq experiment to compare the global gene expression between native levels of *pcdR* and *pcdR* overexpression. To avoid severe cell lysis, the engineered *pcdR* overexpression strain was grown in shaking media to early exponential phase. Cultures were then split into two; in one, IPTG was added at a final concentration of 25 μM, and the other without IPTG. Cells were harvested 15 minutes after the addition of IPTG to induce *pcdR* expression. At this time point, the cell density was still at the tipping point to decline, but it was not dropping yet (Figure 6A).

Results of RNA-seq show that there are about 180 differentially expressed genes (among a total of 4238 genes analyzed) under with and without *yhjH* overexpression conditions when we applied a log2 fold change of ±1 as the cutoff (Figure 7). Interestingly, almost the entire set of SPβ prophage genes were significantly upregulated upon *pcdR* overexpression. In locus order, genes in the entire SPβ genome are upregulated with a log 2-fold change ranging from 0.84 to 2.48, with most genes falling above a log 2-fold change of 1 (Table 1). Another interesting gene that was found is *iseA*, encoding an autolysin inhibitor that interacts with the D-L-endopeptidase domain of major autolysins to inhibit their lytic activities (27). The *iseA* gene was downregulated when *pcdR is* overexpressed (log2 fold change of -2.02).

**Figure 7:**
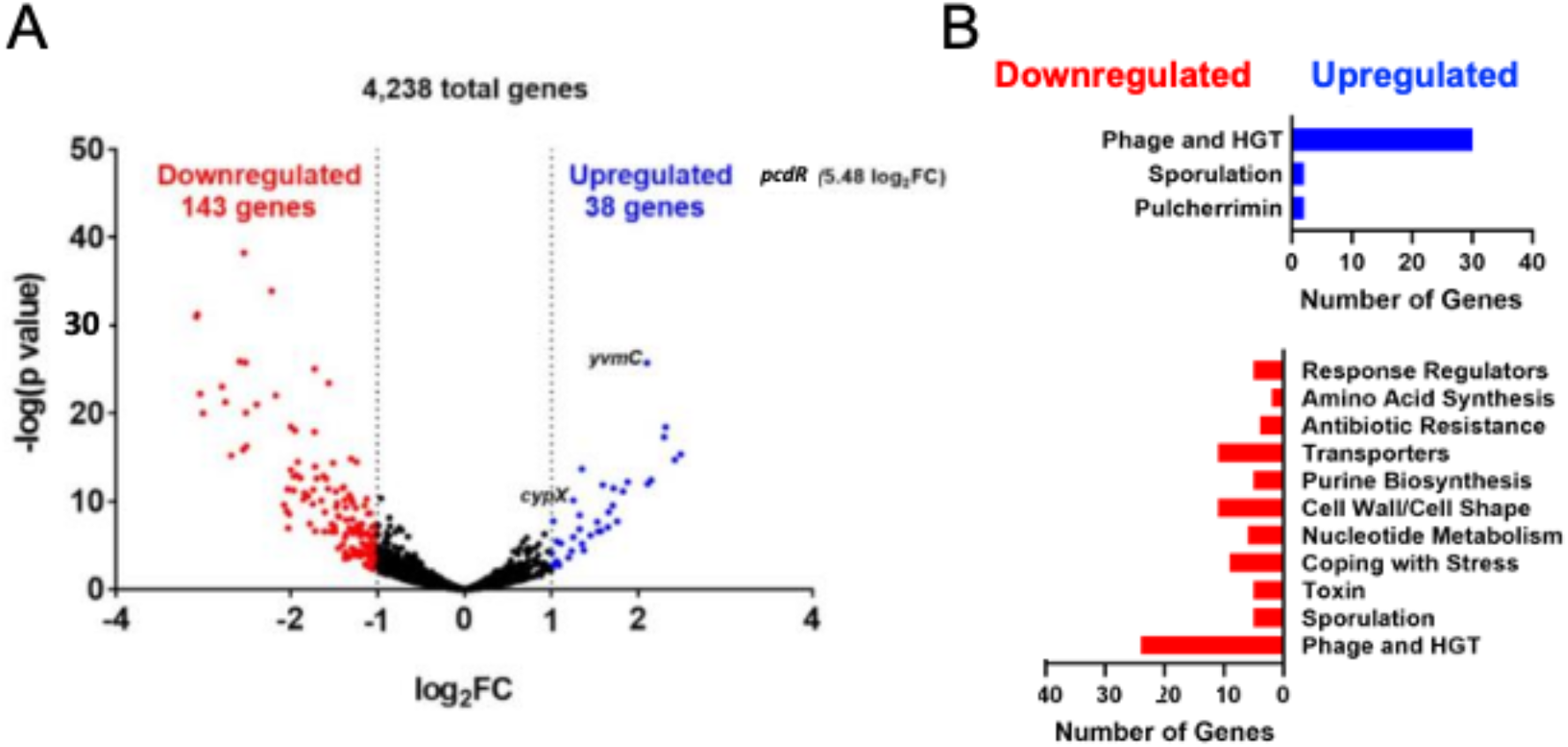
RNA-seq results comparing expression of genes under with and without *pcdR* overexpression conditions. (A) Volcano plat of the RNA-seq results. Based on the log2FC cutoff values of ±1, 38 genes were significantly upregulated (blue dots) while 143 genes (red dots) were significantly downregulated upon *pcdR* overexpression. **(B)** Functional categories of the differentially regulated genes upon *pcdR* overexpression. A large number of the upregulated and downregulated genes belonged to the prophage genes in the *B. subtilis* 3610 genome.

**Table 1:** Induction of SPβ prophage genes (in locus order) upon *pcdR* overexpression.

| Gene name | log2FC | Description |
| --- | --- | --- |
| *yodR* | 1.0866 | similar to butyrate-acetoacetate CoA-transferase |
| *yotB* | 0.8661 | unknown |
| *yosX* | 0.9596 | unknown |
| *yosH* | 0.8990 | unknown |
| *yosF* | 0.8809 | unknown |
| *yosE* | 0.9239 | unknown |
| *yosD* | 1.0103 | unknown |
| *yosC* | 1.0404 | unknown |
| *yosB* | 0.9895 | unknown |
| *yorW* | 0.8874 | unknown |
| *mtbP* | 0.9166 | DNA methyltransferase, restriction modification system |
| *yorM* | 1.3235 | unknown |
| *yorI* | 1.3250 | similar to putative replicative DNA helicase |
| *yorH* | 1.3513 | unknown |
| *yorG* | 1.6585 | unknown |
| *yorF* | 1.7139 | unknown |
| *yorE* | 1.2510 | unknown |
| *yoqR* | 1.1959 | unknown |
| *yoqP* | 1.0769 | unknown |
| *yoqL* | 1.0619 | unknown |
| *yoqK* | 1.5682 | unknown |
| *yoqJ* | 1.5263 | unknown |
| *yoqI* | 1.3709 | unknown |
| *yoqH* | 1.5921 | unknown |
| *yoqG* | 1.4500 | unknown |
| *yoqF* | 1.7043 | unknown |
| *yoqE* | 1.6504 | unknown |
| *yoqD* | 1.8744 | similar to phage-related DNA-binding protein anti-repressor |
| *yoqC* | 1.8218 | unknown |
| *yoqB* | 2.1439 | unknown |
| *yoqA* | 2.3122 | unknown |
| *yopZ* | 2.1044 | unknown |
| *yopY* | 2.2947 | unknown |
| *yopX* | 2.4832 | unknown |
| *yopW* | 2.4166 | unknown |
| *yopV* | 1.5351 | unknown |
| *yopP* | 0.8465 | unknown |
| *yopO* | 1.0045 | similar to transcription regulator (Xre family) |
| *yopN* | 0.8691 | control of lysis/lysogeny in the SP beta phage |
| *aimR* | 0.8396 | phage lytic/lysogenic transcription regulator |
| *yonX* | 1.7563 | unknown |
| *yonV* | 1.1092 | unknown |

### An in-silico DNA binding motif search for PcdR

MarR-type repressors typically bind 16 to 20 bp short inverted repeat sequences slightly upstream of their promoters (26). A repeat sequence matching these conditions was identified both in front of the *yhjG-pcdR* operon and in the intergenic region between *yhjG* and *pcdR* (Figure 8A-C). We constructed a strain with four base-pair changes within that sequence to confirm that repression by PcdR is specific to the proposed binding motif (Figure 8D). There was significantly more promoter activation of the mutated promoter, indicating a loss of repression due to the point mutations within the binding motif (Figure 8E). This result supports the hypothesis that PcdR represses the genes by binding to the predicted binding motif.

**Figure 8:**
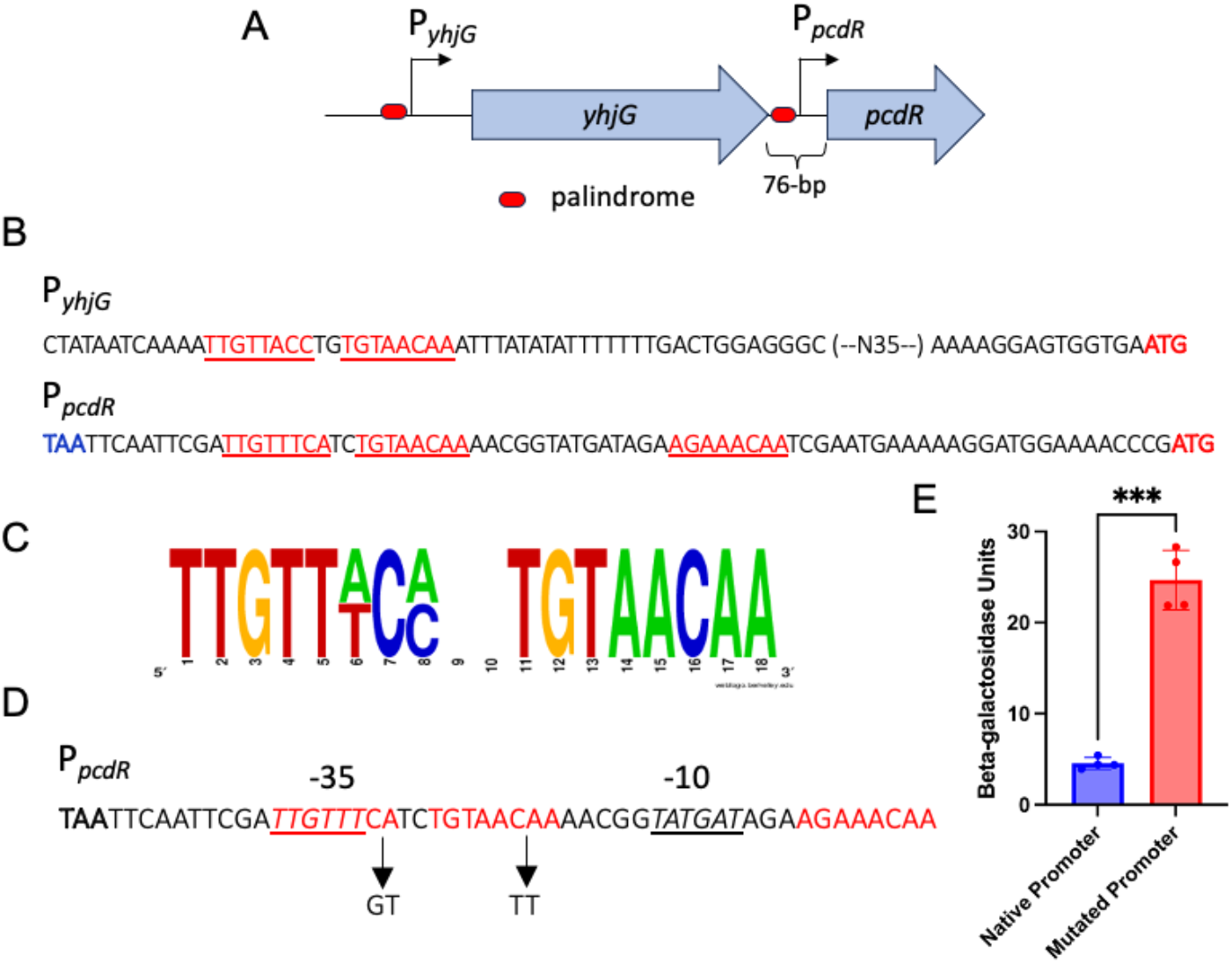
Analysis of putative PcdR binding motif and site-directed mutagenesis. (A) An illustration of the *yhjG-pcdR* operon. Short palindromic sequences overlap with the predicted promoter sequences upstream of both *yhjG* and *pcdR*. The intergenic region between *yhjG* and *pcdR* is 76-bp in size. **(B)** DNA sequences of the promoters of *yhjG* and *pcdR*. Underlined are the palindrome sequences in the promoters of *yhjG* (upper) and *pcdR* (lower) predicted to recognized by PcdR. **(C)** Putative binding motif of PcdR. Constructed from putative binding motifs present upstream of *yhjG* and *pcdR* with the tool provided at https://weblogo.berkeley.edu/. **(D)** Site-directed mutagenesis of the putative PcdR binding motif in the intergenic promoter of *pcdR*. Underlined are the predicted –10 and –35 sites of the *pcdR* internal promoter. The indicated point mutations were made within the putative binding motif (in red). **(E)** Activities of the mutated vs. native promoter of P*pcdR*-*lacZ* was tested in a wild-type strain background. Cultures were harvested after 3 days of incubation in pellicle biofilm conditions. Cells were mildly sonicated and assays of β-galactosidase activities were conducted. Assays were done in triplicate. *** represents P value < 0.0002.

An in-silico search for the PcdR DNA binding motif in the *B. subtilis* genome was conducted. The search tool provided in the website (http://genolist.pasteur.fr/SubtiList/) was used to identify all places in the genome matching that sequence and fitting a few parameters. It was specified that the sequence would need to have between zero and four base pairs between the two halves, that there could be one or two mismatches per side, and that it would need to fall less than 300 base pairs before a gene and not fall within another gene. More than a dozen genes were found as hits in this binding motif search, including the *iseA* gene (Table 2).

**Table 2:** Genes identified in in-silico PcdR DNA binding motif search.

| *Gene* | *Number of mismatches* | *Distance from the gene* | *RNA-seq log2 FC* | *Function* |
| --- | --- | --- | --- | --- |
| *pcdR* | 1 | -66 | 5.49 | unknown |
| *yhjG* | 1 | -94 | -0.45 | unknown; similar to monooxygenase |
| *ylqF* | 1 | -122 | 0.04 | unknown |
| *kapD* | 1 | -483 | -0.25 | inhibitor of the KinA pathway to sporulation |
| *iseA* | 2 | -50 | -2.02 | protein inhibitor of autolysins |
| *yabD* | 2 | -65 | N/A | unknown |
| *nasD* | 2 | -86 | -0.35 | assimilatory nitrite reductase |
| *oppA* | 2 | -137 | 0.03 | oligopeptide ABC transporter |
| *pucJ* | 2 | -141 | 0.00 | pucJ |
| *ybbB* | 2 | -176 | -0.37 | unknown |
| *yoeA* | 2 | -253 | -0.24 | unknown |
| *fliS* | 2 | -320 | 0.58 | flagellar protein |

## Discussion

PcdR is an uncharacterized, putative MarR-type transcription regulator in *B. subtilis*. PcdR was first implicated as relevant to biofilm development through the RNA-seq comparing global gene expression in the presence and absence of the biofilm-associated pigment pulcherrimin. The *pcdR* gene is downregulated in the absence of pulcherrimin. In this work, we further characterized the transcriptional regulation of the *pcdR* gene. The gene is positively regulated by the presence of PA and negatively regulated by both PchR, the repressor for the PA biosynthetic genes, and PcdR itself. Importantly, we uncovered a novel cell death phenotype triggered by *pcdR* overexpression.

The role of programmed cell death (PCD) during biofilm development has been studied in model bacteria such as *Staphylococcus aureus*. Regulated cell death can be used to eliminate damaged cells from a population, to contribute to the release of extracellular proteins and DNA, or to promote 3D structures in a biofilm (18, 19). In *B. subtilis*, areas of high cell death were observed to co-localize with elevated wrinkles in the biofilm architecture. Localized cell death also contributes to biofilm assembly and fruiting body formation (28). Nevertheless, the mechanistic cause of localized cell death in *B. subtilis* biofilms is still unclear. It is possible that control of regulated cell death during biofilm development in *B. subtilis* involves multiple pathways. We propose that PcdR and PcdR-regulated genes play an important role in inducing regulated cell death during biofilm formation.

Two interesting directions for the mechanism of this cell death phenotype were raised from the RNA-seq of PcdR overexpression conditions and the in-silico PcdR binding motif search. Firstly, a significant fraction of SPβ prophage genes are upregulated under PcdR overexpression conditions, and SPβ repressors are significantly downregulated. In locus order, SPβ prophage genes are upregulated at a log 2-fold change between 0.84 and 2.48, where any change above 1 is considered significant. Considering that we only allowed *yhjH* induction for 15 min before cell harvest (to avoid severe cell lysis), the experimental setting could have limited the fold changes in expression of YhjH-regulated genes.

The SPβ phage is a temperate prophage in *Bacillus* that is inserted into sporulation genes (29). The SPβ prophage is excised from the sporulation gene locus where it is inserted to yield a functional *spsM* gene and permit progression in spore formation (30). It is not clear whether the SPβ phage is activated into a lytic cycle during *B. subtilis* biofilm formation. Nevertheless, the lysis associated with *pcdR* overexpression could be caused by the bacterial cell hijacking a phage lytic gene or pathway, rather than full induction of the phage lytic cycle.

Prophages and prophage genes have been shown to play roles in horizontal gene transfer, bacterial evolution, biofilm development, and programmed cell death (17, 31). In *Bacillus anthracis*, it was shown that phage-encoded sigma factors are necessary to induce expression of biofilm genes. These phages co-evolve with their hosts to transfer genes necessary for life in different environments, like the worm gut or soil ecosystem (32). Prophages have been shown to contribute to biofilm formation via induction of programmed cell death in *Staphylococcus pneumoniae* and *Pseudomonas aeruginosa* (21, 33). In the study of *S. pneumoniae*, biofilm development was diminished in an autolysin mutant and a phage mutant, but was completely abolished in the double mutant, indicating that both phage and autolysin activity contribute to the programmed cell death necessary to provide eDNA and proteins to the biofilm matrix (33). In the late stages of the biofilm life cycle, cells sporulate, lyse, and enter a planktonic state to disperse into the environment (34). This stage of biofilm development has shown to be mediated by phage in *P. aeruginosa* (35). Prophages are not simply parasites to bacteria; they contribute in many ways to bacterial development. The results of this study implicate the SPβ phage as a possible mechanism to induce programmed cell death to support biofilm development of *B. subtilis*.

The other potential cause of lysis raised by RNA-seq and the PcdR DNA binding motif search was the gene *iseA*. IseA is a secreted protein that inhibits the function of the DL-endopeptidase LytE, LytF, CwlC, and potentially others by mimicking their substrate (27). These DL-endopeptidases are autolysins, which degrade the cell wall for various reasons. Some autolysins degrade the cell wall create space for cell wall elongation or to achieve cell separation, but some have been shown to induce lysis for programmed cell death (19). In the *pcdR* overexpression RNA-seq, it was found that the *iseA* gene was downregulated with a log 2-fold change of -2.02. Furthermore, a putative PcdR binding motif was identified in the promoter region of the *iseA* gene, located just 50 base pairs upstream of the coding sequence. It has been shown that the *iseA* mutant exhibits cell lysis upon nutrient depletion and is induced by cell envelope-targeting antibiotics (36). The induction of *iseA* in the presence of antibiotics inhibits autolysin activity to slow cell wall turnover, thus protecting the cell from these cell wall-acting antibiotics. MarR-type transcription repressors are commonly modulated by the presence of antibiotics which bind these transcription factors as ligands (26). The binding of ligand alters their shape, causing derepression. Thus, PcdR may also function in vivo to derepress transcription of *iseA* in the presence of antibiotics. To more clearly characterize the regulon of PcdR, a ChIP-seq experiment could be completed in the future to determine which precise DNA sequences are bound by PcdR.

Bacteria use quorum-sensing systems to coordinate behaviors in multicellular systems. One class of understudied signaling molecules in bacteria is cyclodipeptides (14). These molecules cyclize around the amino acid backbone of peptide residues via the action of cyclodipeptide synthases or other non-ribosomal peptide synthases (NRPSs). Cyclodipeptides have been recently identified to have a number of signaling functions in the gut microbiome and biofilms. It is important to note that PA is a modified cyclodipeptide and has never before been indicated as a signaling molecule. We hypothesize that PA could be modulating the activity of PchR, thus controlling the death phenotype effected through *pcdR* expression. Future studies of the signaling capabilities of PA are needed to elucidate how the pigment can affect changes in genetic expression both of its own repressor, PchR, and downstream genes like *pcdR*.

PcdR may be induced by the activity of pulcherrimin to contribute to homeostasis in a biofilm setting. PcdR is a novel transcription factor with important roles in programmed cell death in *B. subtilis* biofilms. Due to this exploration, we have renamed YhjH to PcdR, derived from the phrase “pulcherriminic acid cell death regulator”.

## Methods

### Media and Materials

Strains used in this study are listed in Table 3. *Bacillus subtilis* strain NCIB 3610 is designated as the wild type strain in this study. All other *B. subtilis* strains used in this study are derivatives of 3610 with various genetic modifications. *B. subtilis* PY79 and *Escherichia coli* DH5α were utilized to clone DNA constructs across strains. All strains were cultured at 37°C in lysogenic broth (LB) (10g tryptone, 5g yeast extract, and 5g NaCl added to enough water for liter of broth). Biofilm assays utilized the biofilm-inducing medium LBGM (1% (v/v) glycerol and 100 µM MnSO_4_ in LB broth). Solid agar media was created by adding 15 grams of agar to 1000 mL broth (1.5% w/v). Antibiotics were added to medium as needed at the following concentrations for *B. subtilis*: 200 µg/mL spectinomycin, 10 µg/mL tetracycline, 7 µg/mL chloramphenicol, and 1X macrolide-lincosamide-streptogramin B (MLS). *E. coli* was cultured with 100 µg/mL ampicillin.

**Table 3:** Strains used in this study.

| Strain | Description |
| --- | --- |
| LA003 | Wild-type 3610 |
| LA010 | $\Delta yvmC\text{-}cypX::tet^R$ |
| LA016 | DH5α strain for molecular cloning |
| LA383 | $\Delta pcdR::tet^R$ |
| LA406 | $\Delta pchR::erm^R$ |
| LA606 | $\Delta pchR::erm^R \Delta pcdR::tet^R$ |
| LA354 | $amyE::P_{yhjG}\text{-}gfp, spec^R$ |
| LA401 | $\Delta pcdR::tet amyE::P_{yhjG}\text{-}gfp, spec^R$ |
| LA355 | $\Delta yvmC::tet amyE::P_{yhjG}\text{-}gfp, spec^R$ |
| LA408 | $\Delta yvmC\text{-}cypX::tet^R, amyE::P_{hyspank}\text{-}yvmC\text{-}cypX, thrC::P_{yhjG}\text{-}gfp, spec^R, erm^R$ |
| LA422 | $\Delta pchR::erm^R amyE::P_{yhjG}\text{-}gfp, spec^R$ |
| LA642 | $amyE::P_{pcdR}\text{-}lacZ, cm^R$ |
| LA644 | $\Delta pcdR::tet^R amyE::P_{pcdR}\text{-}lacZ, cm^R$ |
| LA448 | $amyE::P_{hpspank}\text{-}pcdR, spec^R$ |
| LA465 | $amyE::P_{hpspank}\text{-}pcdR$ (V107G), $spec^R$, suppressor 8 |
| LA680 | $amyE::P_{pcdR}\text{-}lacZ$ mutated binding site, $cm^R$ |

### Polymerase Chain Reaction (PCR)

For all PCRs used in this study, a 2X OneTaq mastermix (New England Biolabs) was mixed with 1 µL template DNA and a final concentration of 0.5 µM of each primer.

### Strain Construction

To construct a *pcdR* gene deletion, the method of long-flanking homology (LFH) was applied (37). To do so, two ∼1000-bp regions on either side of the *pcdR* coding region in the *B. subtilis* genome were amplified by PCR. The PCR products, after gel purification (Qiagen), were then used as megaprimers to amplify a region of DNA with the flanking regions surrounding an antibiotic resistance cassette. This entire product was then transformed into *B. subtilis* PY79. The transformants were selected for double crossover recombination in the *pcdR* locus in PY79. Deletion of the *pcdR* gene on the chromosomal locus was confirmed by PCR and DNA sequencing. The genomic DNA was then isolated from the above verified transformant and introduced into *B. subtilis* 3610. The *pcdR* gene deletion in 3610 was again verified by PCR and DNA sequencing. All oligonucleotides used in this study are listed in Table 4.

**Table 4:**
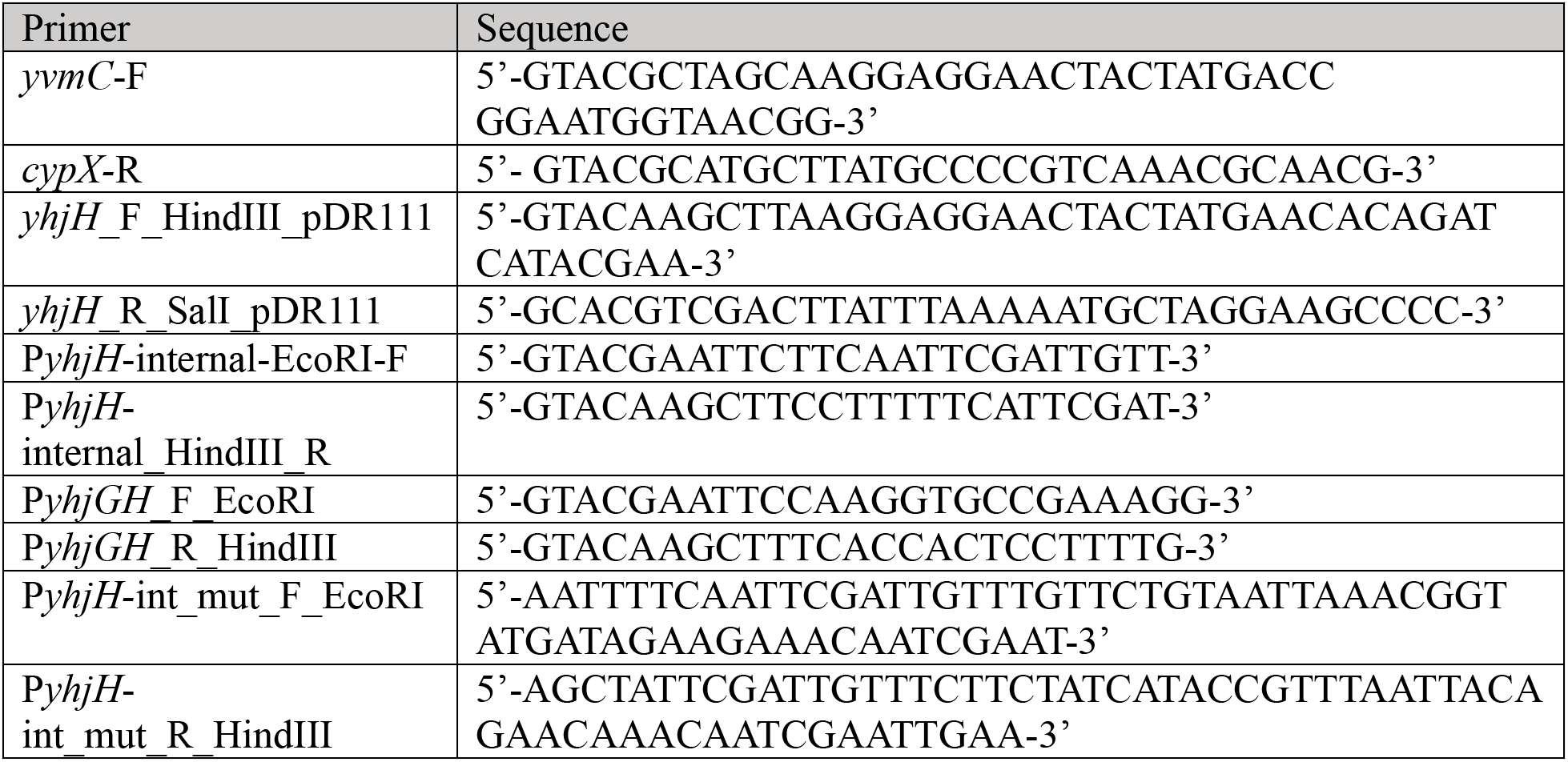
Oligonucleotides used in this study.

To build promoter fusion strains, the promoter region of the target gene was amplified via PCR from wild-type genomic DNA. In this PCR step, primers would incorporate restriction enzyme sequences on either end of the target DNA. Sequences would be chosen to match those in specified locations in certain plasmids. Next, plasmids and PCR products would be digested with the corresponding restriction enzymes and ligated together with T4 DNA ligase (New England Biolabs). The ligated plasmid would be transformed into *E. coli* using a heat-shock transformation protocol, then *E. coli* strains were grown and miniprepped to collect the recombinant plasmid. This plasmid was then transformed directly into *B. subtilis* 3610. The same process was used to build the *pcdR* overexpression strain, but the target DNA was *pcdR* itself.

### RNA-seq

For RNA-seq comparing global transcriptomes with and without overexpression of *pcdR*, the engineered strain (LA448, *amyE*::P*_hyspank_*-*pcdR*) was cultured overnight for 16 hours. Then the overnight culture was diluted 100-fold into fresh media and grown until the OD600 of the culture reaches 0.5. The culture was then split into 2 equal volumes. For one volume, IPTG was added to a final concentration of 25 µM, and the other volume without addition of IPTG. Both cultures were incubated for another 15 minutes. Next, cultures were spun down, resuspended with 1 mL of RNA protect solution (Invitrogen), and incubated for 5 minutes at room temperature. Finally, pellets were flash frozen with liquid nitrogen and sent for RNA-sequencing (Genewiz, NJ, USA).

### Biofilm Development

To develop pellicle or colony biofilms, strains were grown overnight in shaking LB broth. The next morning, fresh LB broth was used to inoculate overnight culture and cultures were grown until cells reached exponential phase. Then, shaking culture was used to inoculate in a 1:1000 dilution to the biofilm-inducing media LBGM. Pellicle Biofilms were then statically incubated at 30°C for indicated periods of time before harvest, typically 2-3 days.

### Beta-galactosidase Assay

To measure promoter activation of cells in biofilms via beta-galactosidase assay, cells were grown until the desired time point and harvested. When cells were harvested, the biofilm culture was sonicated to completely disrupt the biofilm, then the density of the cells was measured via OD600. Cells were then centrifuged, supernatant was removed, and the samples were frozen at - 80°C.

For the beta-galactosidase assay itself, cells were thawed and resuspended in 1 mL Z-buffer (0.06 M Na_2_HPO_4_, 0.04 M NaH_2_PO_4_, 0.01 KCl, 0.001 M MgSO_4_, and 0.27% (v/v) β-mercaptoethanol). 20 mg/ml lysozyme was added to small groups of samples at a time and incubated for 15 minutes at 30°C to allow cell lysis to occur. Next, 200 µL of a solution of 4 mg/mL ortho-nitrophenyl-β-galactoside (ONPG) in Z-buffer was added to start a color changing reaction. 500 µL of 1 M sodium carbonate (Na_2_CO_3_) was added to stop the reaction after a sufficient color change was induced. The time between addition of ONPG and Na_2_CO_3_ was measured.

To quantify the extent of this reaction, samples were spun down for 10 minutes and two wavelengths were measured: OD420 and OD550. β-galactosidase activity was quantified in Miller units with the following formula: [(OD_420_-1.75 ×OD_550_)/(time × volume in mL × OD_600_)] × 1,000.

### Microscopy

Cells were grown to exponential phase in an LB media shaking culture before being spun down, washed in 1x phosphate buffer solution (PBS) and visualized. Cells were imaged with a Leica DFC3000 G camera on a Leica AF6000 microscope. For GFP imaging, the excitation wavelength was set to 450-490 nm and the emission wavelength was set to 500-550 nm.

For time-lapse microscopy, 3-µL cells from the log phase culture were inoculated to the center of an agar pad consisting of LB media with 1.5% agar. IPTG was added to the agar pad at the final concentration of 25 µM. A cover slip was added and cells were imaged using confocal microscopy. Snapshot pictures for the same area of cells were taken every 3 minutes for up to 5 hours.

## Acknowledgements

This work was supported by a National Science Foundation grant (MCB1651732) awarded to Y. Chai. We thank Nicole Cavanaugh and Yinghao He from the Chai lab for their help with guidance and troubleshooting for experiments.

## Conflict of Interest

The authors declare no competing interests.

## Data Availability

The authors declare that all data supporting the findings of this study are available in this article and its Supplementary Information files, or from the corresponding author upon request.

## Author Contributions

G.M. and L.L.A. performed the experiments and strain construction; G.F. performed bioinformatics and data analysis; V.G. provided critical suggestions during manuscript preparation and experimental design. G.M., L.LA., and Y.C. designed the study and wrote the manuscript.

## Notes

### Competing Interest Statement

The authors have declared no competing interest.

