## Supplementary Figure 1 for "Discovery of a novel MarR-type transcriptional regulator that controls cell death in *Bacillus subtilis* biofilms"

```
PchR -MSDLTKQMIYDIYVRLLHLNEQKANTSLQQFFK----EAAEEDVAEIPKNMTSIHVIDC 55
PcdR MNTDHTKRNLFELYAELIHQQEKWEGL-IKAFLSDELRKLDVEHGSKSQLTMTEIHVLSC 59
       :* **: ::::*..*:* :*:   . :: *:.      : *.   :: .**.***:.*

PchR IGQHEPINNAGIARKMNLSKANVTKISTKLIKEEFINSYQLTDNKKEVYFKLTRKGRRIF 115
PcdR VGDNEPINVTSIAEKMNTTKATVSRISTKLLGAGFLHRTQLSDNKKEVYFRLTPAGKKLH 119
     :*::**** :.**.*** :**.*::*****:   *::  **:********:**  *:::.

PchR DLHEKLHKKKELAFYQFLDSFSQEEQKAVLKFLEQLTSTLEAE---QTDGTPDKPVK 169
PcdR SLHKYYHQKAEQRFLSFFDRYTEEEILFAERLFRDLVTKWYPSSEEIEGGLPSI-FK 175
     .**:  *:* *  * .*:* :::** . :::  .:*.:. .       .* *.  .*
```

**Figure S1. Protein sequence alignment between PchR and PcdR, two MarR-type transcription repressors in *B. subtilis*.** These two proteins share about 42% identity in amino acid sequence. V107 in PcdR (highlighted in red) was mutated to Alanine (V107A) in one of suppressor mutants that we isolated (suppressor 8). Overexpression of pcdR (V107A) no longer triggers cell lysis in suppressor 8.
